## Supplementary Figures 1-5 for "Monitoring myelin lipid composition and structure of myelinated fibers reveals a maturation delay in CMT1A"

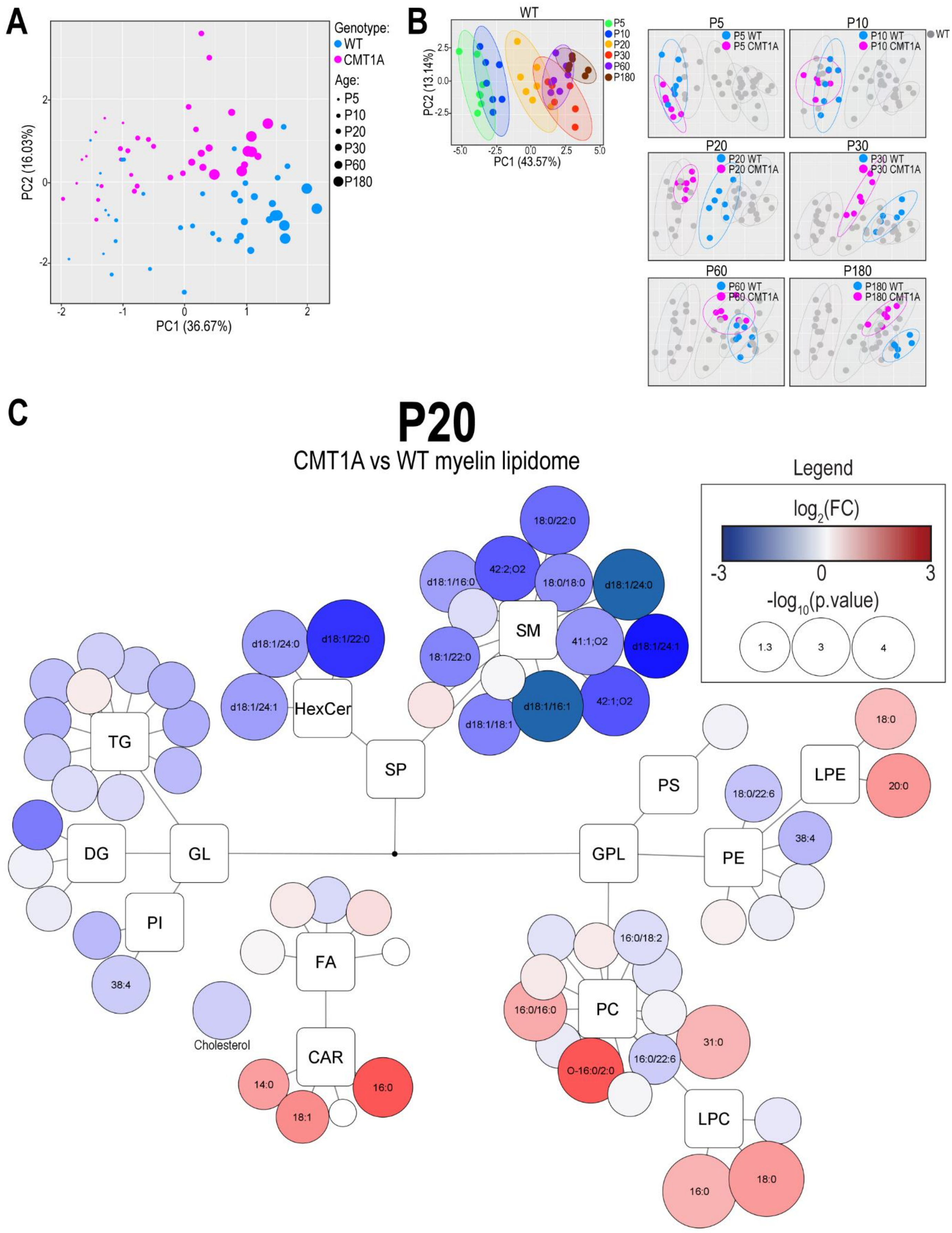


**Supplementary Figure 1. Lipidomics Data**

(A) PCA plot of lipidomic data from WT and CMT1A purified peripheral myelin extracted from the sciatic nerve of P5, P10, P20, P30, P60, and P180 rats. n = at least 6 samples for each group; for each sample, several nerves from different rats were pulled together to obtain enough myelin. (B) PCA plots showing CMT1A samples from each time point together with WT samples, to highlight the maturation delay experienced by CMT1A myelin. (C) Simplified network visualization of lipid metabolism showing alterations in myelin lipid profile of P20 CMT1A rats, as compared to WT. Each dot represents a lipid species, dot size expresses the significance according to p-value, while the color intensity defines the degree of up (red) and downregulation (blue) according to the fold change (as CMT1A vs WT). The lipids showing a statistically significant difference are annotated. n = 7 WT and 7 CMT1A samples; for each sample, sciatic nerves from 5 different animals were pulled together to obtain a sufficient amount of material for the analysis. p.values were calculated using a two-tailed unpaired Student's t-test.

A

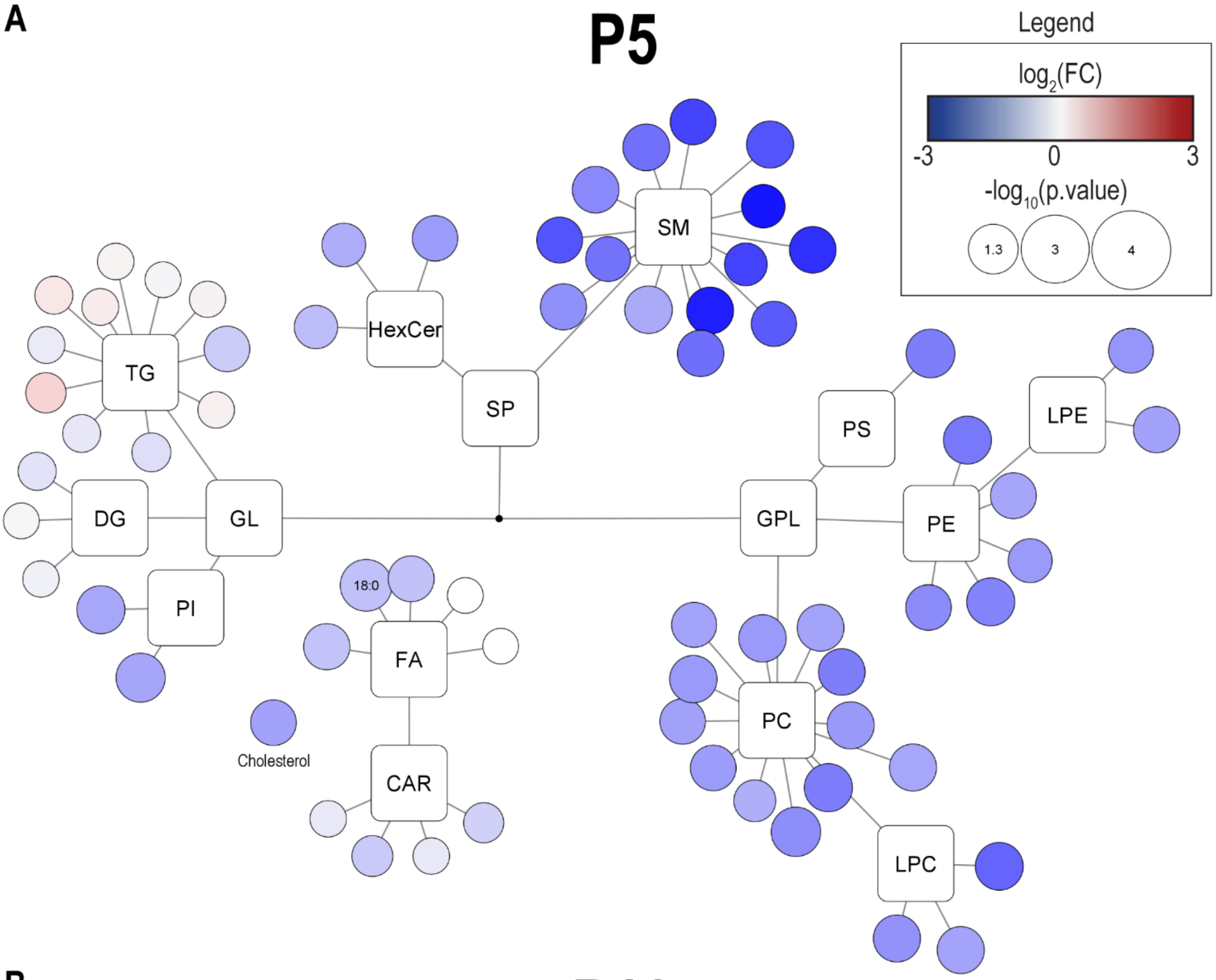
P5
Legend
log2(FC)
-3
0
3
-log10(p.value)
1.3
3
4
SM
HexCer
SP
TG
DG
GL
PI
FA
18:0
CAR
Cholesterol
PS
LPE
GPL
PE
PC
LPC

B

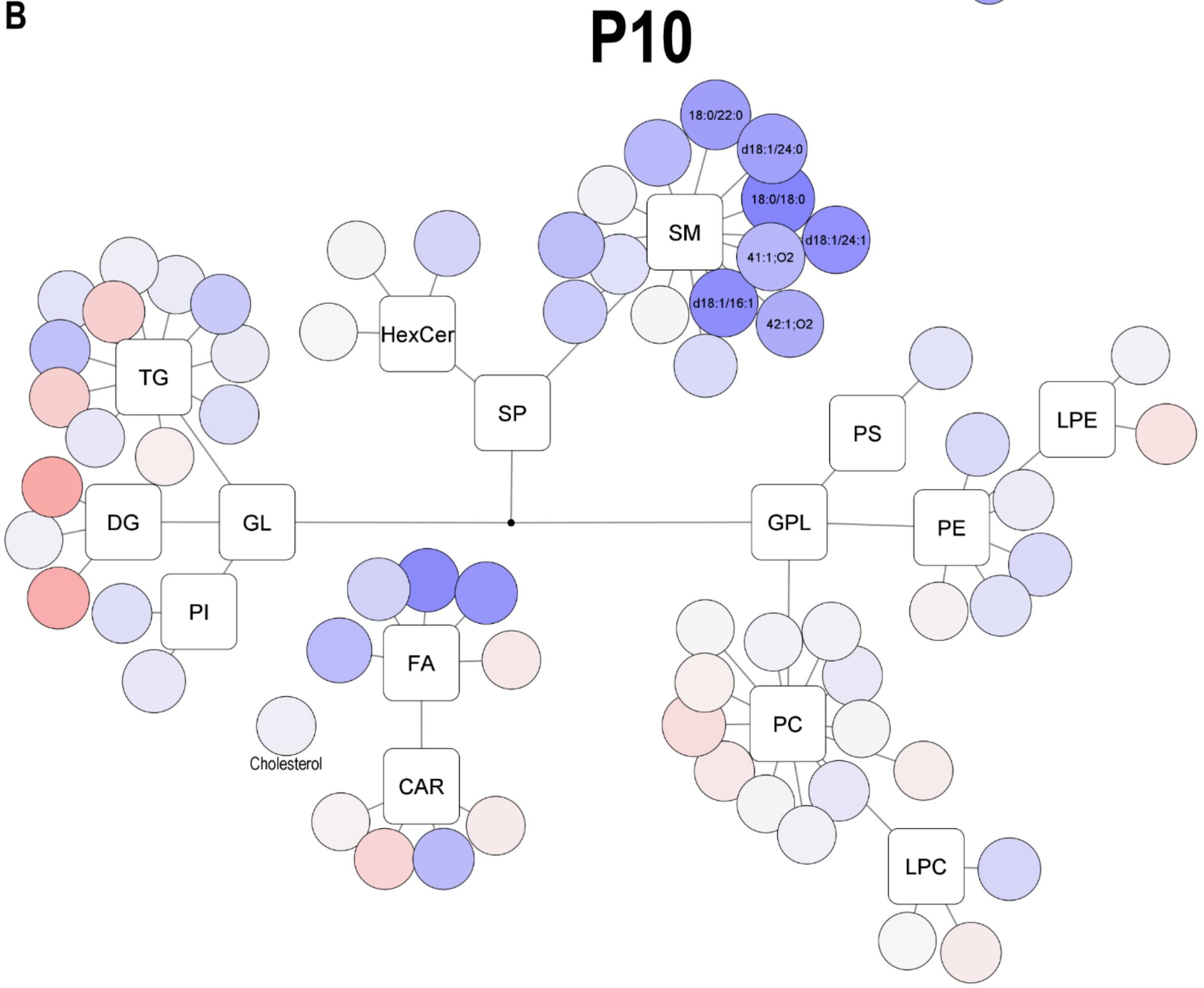
P10
SM
18:0/22:0
d18:1/24:0
18:0/18:0
d18:1/24:1
41:1;O2
d18:1/16:1
42:1;O2
HexCer
SP
TG
DG
GL
PI
FA
CAR
Cholesterol
PS
LPE
GPL
PE
PC
LPC

**Supplementary Figure 2. Lipidomics Data P5 and P10**

Simplified network visualization of lipid metabolism showing alterations in myelin lipid profile of P5 (A) and P10 (B) CMT1A rats, as compared to WT. Each dot represents a lipid species, dot size expresses the significance according to p-value, while the color intensity defines the degree of up (red) and downregulation (blue) according to the fold change (as CMT1A vs WT). The lipids showing a statistically significant difference are annotated. n = 7 WT and 7 CMT1A samples; for each sample, sciatic nerves from 5 different animals were pulled together to obtain a sufficient amount of material for the analysis. p.values were calculated using a two-tailed unpaired Student's t-test.

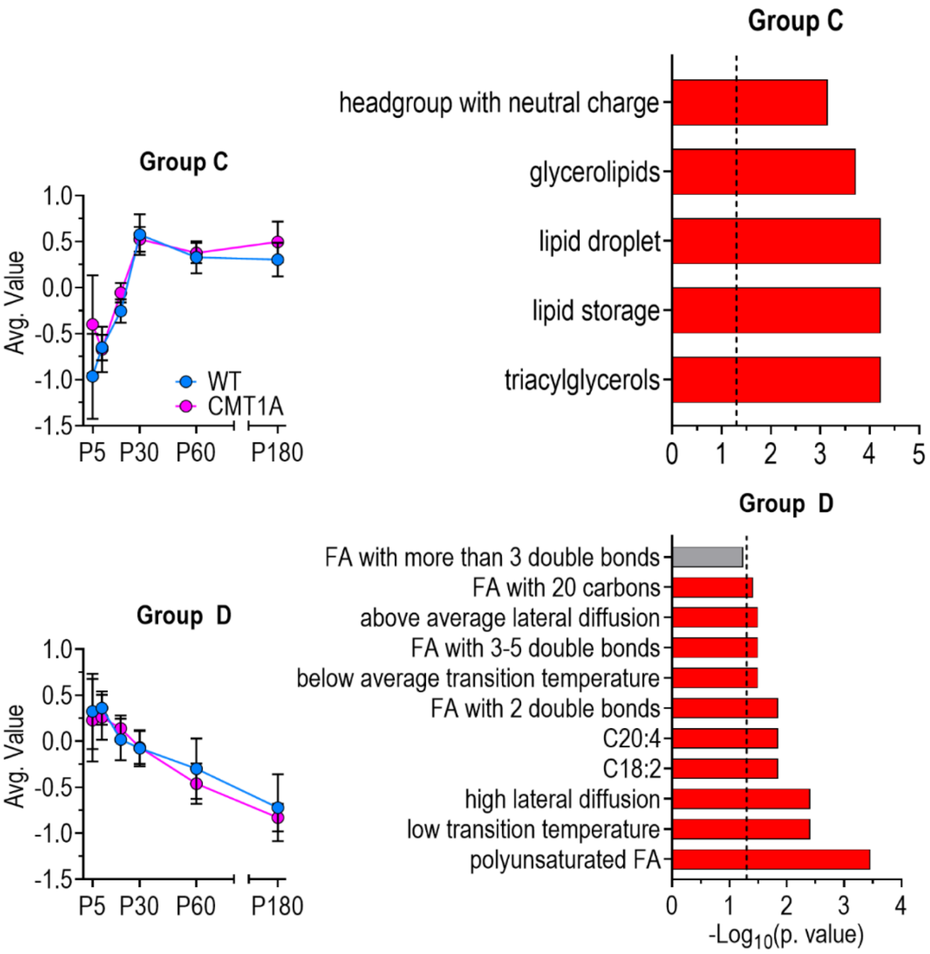


**Supplementary Figure 3. Lipidomics Data**

(A) On the left, average value of lipids in group C (top) and D (bottom); on the right, results of enrichment analysis performed using LION software on lipids in group C (top) and D (bottom). The dotted line marks p.value significance threshold (0.05).

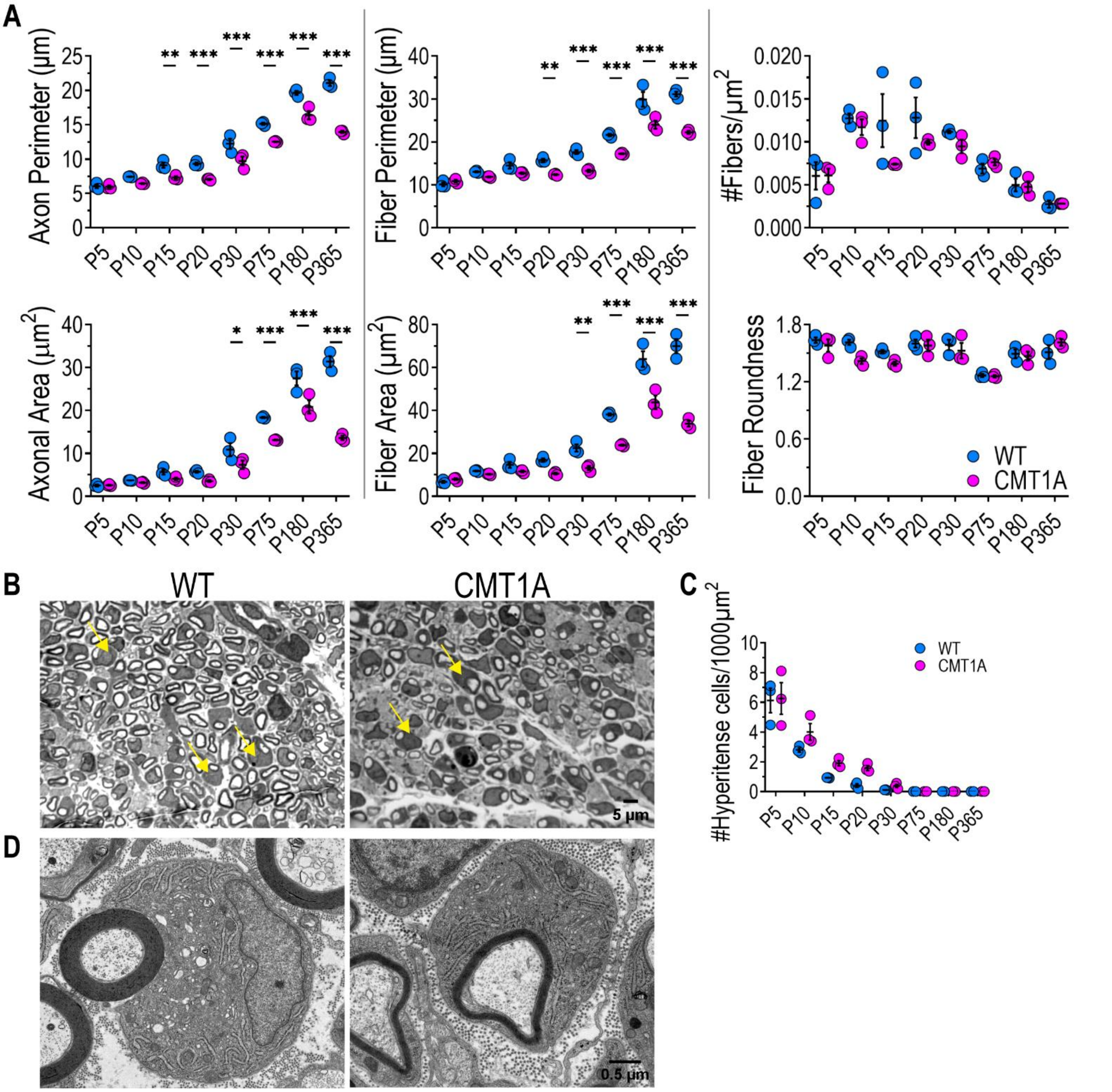


**Supplementary Figure 4. Morphometric Parameters and Hyperintense cells.**

A) Morphometric parameters from WT and CMT1A sciatic nerve sections. (B) Micrographs of cross-sections of P5 WT and CMT1A sciatic nerves stained with toluidine blue; yellow arrows indicate hyperintense cells. (C) Hyperintense cell density in WT and CMT1A sciatic nerves at different time points. n = 3 rats for each genotype and each time point. At P5, at least 800 myelinated fibers were evaluated for each rat; at later time points, at least 3000 fibers were evaluated for each rat. (D) Transmission electron microscopy micrographs of representative hyperintense cells. Scale bar: 0.5 µm. Values are presented as mean ± SEM. * = p.value < 0.05, ** = p.value < 0.01, *** = p.value < 0.001. p.values were calculated using Kruskal–Wallis test followed by Sidak's multiple comparisons tests between the two genotypes for each time point.

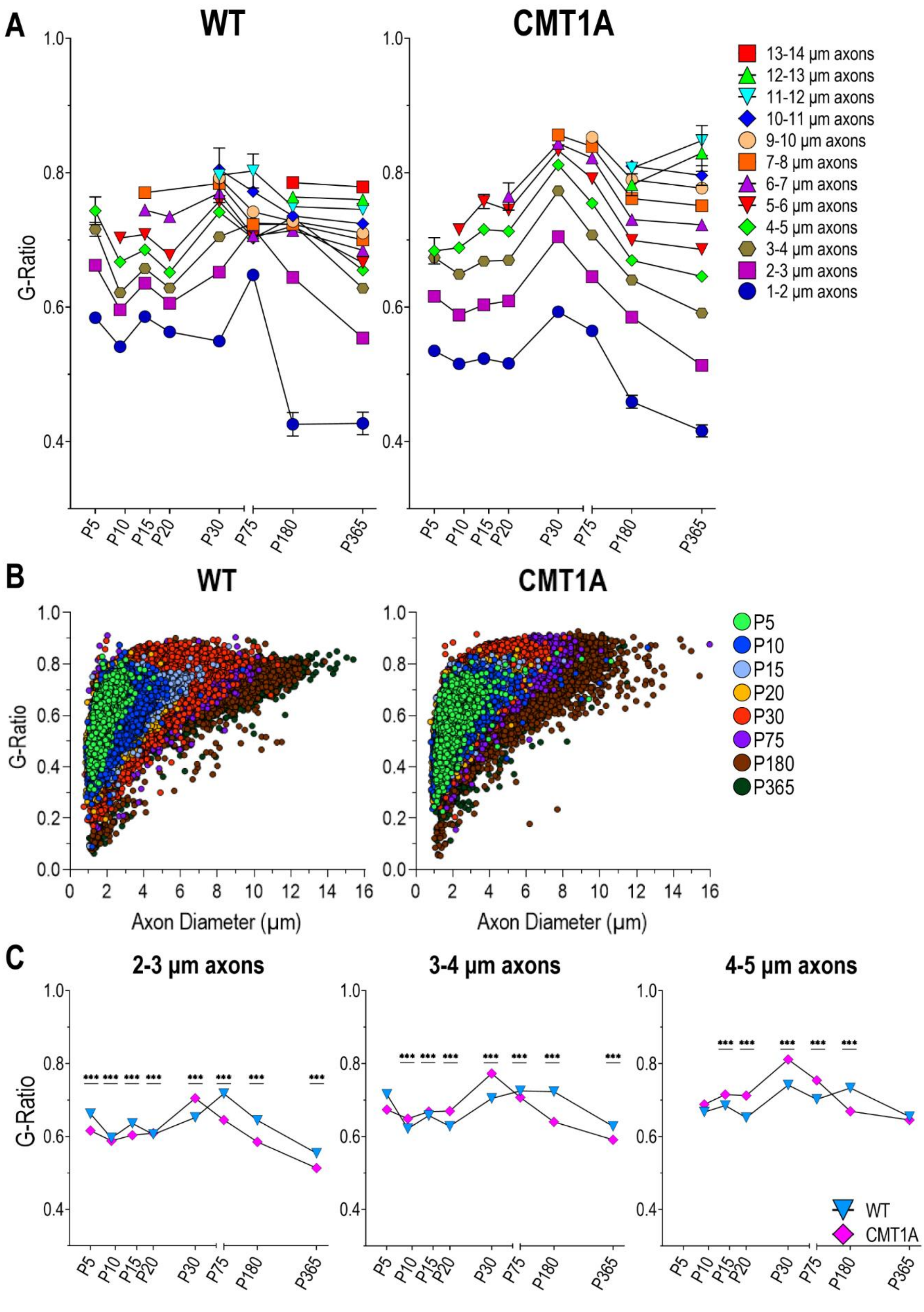


**Supplementary Figure 5. Analysis of G-Ratio evolution during development.**

(A) Developmental trajectories of G-Ratio in WT (left) and CMT1A (right) rat sciatic nerves from early postnatal days to adulthood, measured for each axon class. Fibers were classified based on axon diameter. (B) Scatterplots showing the G-Ratio plotted against the axon diameter of WT and CMT1A sciatic nerve fibers at different time points. (C) Same data as in A, but WT and CMT1A of the same axonal class are plotted together. Values are presented as mean ± SEM; for some values, error bars are not visible because they are covered by the symbol. *** = p.value < 0.001. n = 3000 myelinated fibers from 3 different animals per genotype were evaluated at P5, 9000 myelinated fibers from 3 different animals per genotype were evaluated at later time points. p.values were calculated as in Fig. 3C.

### Supplementary Materials and Methods

#### Quantitative fiber morphology

Fiber Roundness was defined as π × diameter/perimeter. This parameter can be used as a parameter of tissue integrity since it is sensitive to physiological strains acting on myelinated fibers during development; for the same reason, it is a useful criterion to exclude from the analyses fibers displaying artifacts due to tissue processing.

#### Hyperintense cells

"Hyperintense" cells were counted at each time point for the two genotypes. For transmission electron microscopy analysis, 70 nm ultra-thin sections were cut and stained with 1% uranyl acetate and lead citrate solution. Images were collected with a Jeol JEM 1011 (Jeol, Japan) electron microscope, operating with a maximum acceleration voltage of 100 kV, and recorded with a 2 Mp charge-coupled device camera (Gatan Orius SC100).

#### Data visualization

Network visualization of lipid metabolism was generated using Cytoscape (v. 3.10.1) (https://cytoscape.org/).
